# Self-organization and culture of Mesenchymal Stromal Cell spheroids in acoustic levitation

**DOI:** 10.1101/2020.11.16.385047

**Authors:** Nathan Jeger-Madiot, Lousineh Arakelian, Niclas Setterblad, Patrick Bruneval, Mauricio Hoyos, Jérôme Larghero, Jean-Luc Aider

**Affiliations:** Laboratoire de Physique et Mécanique des Milieux Hétérogènes (PMMH), UMR 7636 CNRS, ESPCI Paris, PSL, Paris Sciences et Lettres University, Sorbonne Université, Université de Paris 1, Paris, 75005, France; Unité de Thérapie Cellulaire, APHP, Hôpital Saint-Louis, 1 avenue Claude Vellefaux, F-75010 Paris, Université de Paris, Inserm U976 et CIC de Biothérapies CBT501, F-75006 Paris, France; Technological Core facility of the Institut de Recherche Saint-Louis, Université Paris-Diderot and Inserm, Hôpital Saint-Louis, Paris, France; INSERM U970-PARCC, Paris, France

**Author notes:** these authors contributed equally (Nathan Jeger-Madiot, Lousineh Arakelian).

**Keywords:** Multi-trap acoustofluidic device, Cell culture in acoustic levitation, Scaffold-free 3D culture, Mesenchymal Stromal Cells, Spheroids

## Abstract

In recent years, 3D cell culture models such as spheroid or organoid technologies have known important developments. Many studies have shown that 3D cultures exhibit better biomimetic properties compared to 2D cultures. These properties are important for *in-vitro* modeling systems, as well as for *in-vivo* cell therapies and tissue engineering approaches. A reliable use of 3D cellular models still requires standardized protocols with well-controlled and reproducible parameters. To address this challenge, a robust and scaffold-free approach is proposed, which relies on multi-trap acoustic levitation. This technology is successfully applied to Mesenchymal Stromal Cells (MSCs) maintained in acoustic levitation over a 24-hour period. During the culture, MSCs spontaneously self-organized from cell sheets to cell spheroids with a characteristic time of about ten hours. Each acoustofluidic chip could contain up to 30 spheroids in acoustic levitation and four chips could be ran in parallel, leading to the production of 120 spheroids per experiment. Various biological characterizations showed that the cells inside the spheroids were viable, maintained the expression of their cell surface markers and had a higher differentiation capacity compared to standard 2D culture conditions. These results open the path to long-time cell culture in acoustic levitation of cell sheets or spheroids for any type of cells.

## Introduction

3D cell cultures have shown closer biological properties to cells in a physiological context than 2D cell cultures [1, 2]. Therefore they have become one of the main research and development topics in cell biology of recent years [3]. Nevertheless, the clinical translation of these models has been very limited due to a lack of reproducibility of the results [4]. Most protocols rely on the use of poorly defined hydrogels which affect cells with their chemical and biomechanical properties [5].

Alternatively, several scaffold-free methods, such as hanging drop [6], non-adhesive surface [7] or rotating bioreactor [8], have shown spheroid fabrication as 3D cell culture models. However, some disadvantages such as variation in spheroid size and shape, or lack of cell-matrix interactions still remain unsolved [9].

Thanks to its excellent biocompatibility and versatility, acoustic approaches are promising tools for contactless manipulation of cells [10]. Besides the various applications of cell focusing or separation [11], acoustic methods have been used for tissue engineering and *in vitro* techniques [12]. Many works [13, 14] studied cell behavior with a short-time ultrasound exposure, classically around one hour and used it to instantaneously shape complex geometries like layer [15], multi-layer [16] or spheroid [17] in order to culture it in usual way.

Here, we report the fabrication and the culture of several spheroids over a 24-hour period in an acoustofluidic chip with a well-controlled cell culture environment. The acoustic forces act on the suspended cells [18] by clustering and trapping them in multiple disk-like layers in acoustic levitation, located at the different pressure nodes of a resonant cylindrical cavity [19]. After a self-organization step, the spheroids could be maintained and cultured in levitation, with a system of perfused culture medium.

We designed an acoustofluidic chip made of a PDMS (Polydimethylsiloxane) body bonded to a microscope cover-glass. We chose PDMS material because it is highly biocompatible and permeable to the air, making it ideal for cell culture [20]. It is also transparent, allowing optical accessibility and is easy to process.

A 2 MHz ultrasonic transducer (**Figure 1a**) was inserted into the PDMS body, in direct contact with the fluid, closing the cylindrical cavity. The microscope cover-glass, facing the transducer, acts as an efficient acoustic reflector and allows an optical access to visualize the cell clusters. The geometry of our chip has been optimized in order to increase the number of acoustic traps per chip, up to 30, and to be easily parallelizable. In this study we have used four chips in parallel, leading to the culture of 120 cell spheroids per experiment. To maintain sterility, the chips are single use and disposable.

**Figure 1.**
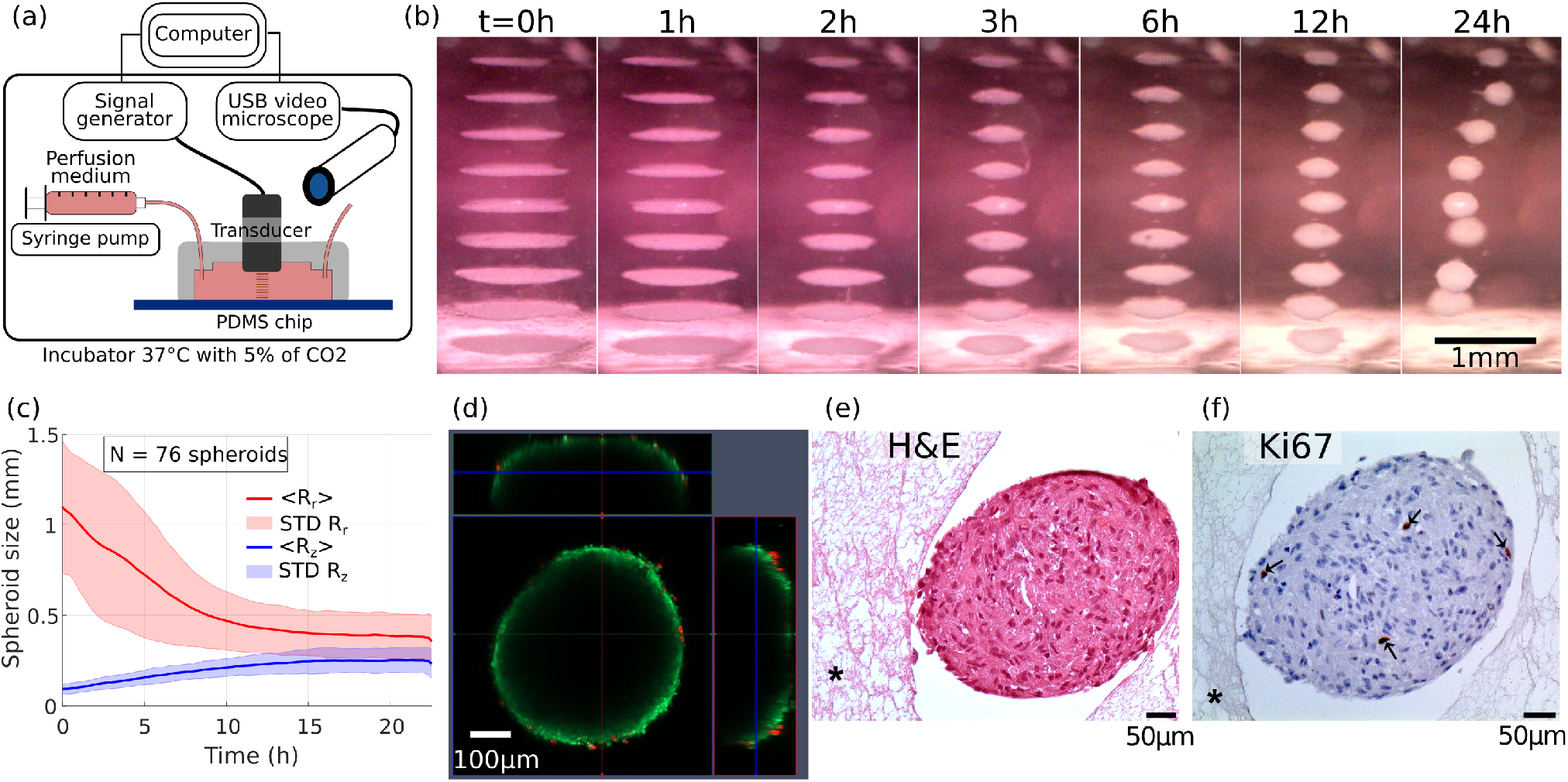
Self-organization of MSCs cells sheets into spheroid shapes and biological viability. (a) Presentation of the experimental setup. A computer drives the wave generation and the acquisition of images. The transducer converts the electrical energy into acoustic energy inside the chip cavity and produced the acoustic levitation of cells over multiple acoustic pressure nodes. A USB digital microscope took side-view pictures and time-lapses of the cell sheets in levitation and self-organization into spheroids. A syringe pump allows a continuous perfusion of the cell medium during the 24h acoustic manipulation. The whole setup, apart from the computer and the generator, was put in an incubator for a well-controlled cell culture environment. (b) Time lapse snapshots of the cell self-organization. The whole time-lapse is available on the movieCellLevitationEx1.mp4. At the beginning, the cells were trapped in monolayers. Then, the cells sheets contracted and the cells self-organized to reach a stable spheroid shape. (c) Time-evolution of the axial *R*_*z*_ and the radial *R*_*r*_ dimensions study of the cells aggregates, averaged over 76 individuals, computed from the time-lapse observations. At *t* = 0h, the widths *R*_*r*_ ranged from 0,75 to 1,5 mm depending on the aggregate location. However, all the heights were similar because of the monolayer shape. After 15h of levitation, every MSC aggregates have reached a stable spheroid shape (*R*_*z*_ ≈ *R*_*r*_). (d) Confocal and fluorescence imagery of a spheroid just after the acoustic levitation. A live/dead kit was used to label the cells. Because of the limited optical penetration or the substrate diffusion, only the cells at the surface were visible and a majority of them showed a living signal. (e) Histology of MSC spheroids. After a 24h levitation, spheroids were collected from the chip and were immediately fixed in parafolmaldehyde 4% overnight. The spheroids were then embedded in fibrin network (described by the symbol * on the images) for an easy handling of paraffin inclusion and histological preparation. Spheroids were stained with H&E for general structure evaluation and (f) Ki67 for proliferation.

We first demonstrated the functionality of our system using a suspension of 10μm polystyrene beads close to the targeted cells dimensions. As soon as the acoustic generator was turned on, the particles were trapped in the acoustic nodes as monolayers and remained stable as long as needed. This result is illustrated for a 90 hours experiment on the video movieBeads.mp4.

We then evaluated our system for culturing human cells. MSCs are primary cells that can been isolated from various tissue sources including the umbilical cords, adipose tissue and the bone marrow. They are characterized by their ability to adhere to plastic, by the expression of certain surface markers, as well as their capacity to differentiate into adipocytes, osteoblasts and chondrocytes. These cells are used in many clinical and tissue engineering applications. We decided to use this cell model as a proof of concept for our system. We isolated these cells from human adipose tissue.

## Results

### 24 hour acoustic levitation of MSCs and self-organization into spheroids

For the acoustic levitation experiments, the cells were injected in the cavity at a concentration of 1.5 million cells/ml and cultured for 24 hours. The setup has been optimized to allow a constant flow of nutrients. The cell culture medium was injected at a rate of 20μL/h, which allowed constant medium renewal without disturbing the cells aggregates in acoustic levitation. The observation of the cells’ collective behavior in levitation showed that they first formed monolayers in the same way as for the beads. Then, gradually, during the first few hours, they spontaneously reorganized and formed spheroids of a typical diameter close to 450μm (Figure 1b, movieCellLevitationEx1.mp4 and movieCellLevitationEx2.mp4). This behavior was simultaneously observed in all the cell layers in acoustic levitation. This illustrated the self-organizational behavior of living cells, which was not observed with polystyrene beads.

### Time evolution of the spheroid sizes

The dynamics of the self-organization into spheroids was studied through image processing of the video time series. The axial *R*_*z*_ and radial *R*_*r*_ dimensions of the spheroids were measured for every spheroid and averaged at each time step. We show on Figure 1c the time evolution of the mean values < *R*_*z*_ > (*t*) and < *R*_*r*_ > (*t*). One can see that after 12 hours, all the cell sheets have turned into stable spheroids elongated along the radial dimension. The average diameter of the spheroids, obtained on 120 samples, is 400μm.

### Viability and proliferation of the MSCs spheroids after the 24 hour acoustic levitation

We evaluated the viability of the cells after a 24 hours culture in acoustic levitation using calcein and propidium iodide, which showed that the majority of the cells were alive at the surface of the spheroid (Figure 1d). However, due to the optical limitation of the confocal microscopy or substrate diffusion, it was not possible to visualize the core of the spheroids with this technique. We therefore evaluated the integrity of the structures by histology. Hematoxylin and eosin (H&E) staining showed that the spheroids were filled structures, with no necrosis in their cores (Figure 1e). Furthermore, the intact form of the cell nuclei indicated that they were not apoptotic. The evaluation of cell proliferation by Ki67 staining showed that most cells were not proliferative but a few cells still stained positive, despite being in levitation (Figure 1f). These results indicate a behavior of the cells inside the spheroids close to that of cells within tissues, which are mainly quiescent while only a few cells proliferate [21].

### Differentiation of the MSCs spheroids

The spheroids were then reseeded in culture plates (**Figure 2a**), and their attachment and spreading were observed with a video microscope (Nikon Biostation, moviePh.mp4 and movieGFP.mp4). The spheroids maintained their potential to re-adhere to plastic and MSCs were shown to proliferate and spread out. In these conditions, MSCs expressed the standard surface markers of mesenchymal cells, thus showing that levitation did not alter these properties (Figure 2b). Furthermore, differentiation into adipocytes after attachment to plastic showed that the cells closest to the center of the spheroid had the highest potential of differentiation and lipid accumulation (Figure 2c). The number of adipocytes observed with the spheroid was much higher than the control condition in 2D culture. When differentiated along the osteoblastic lineage, MSCs cultured in levitation showed higher number and a larger size of osteoblast nodules than in 2D control conditions (Figure 2c), indicating that the physical effects induced by levitation increased the potential of osteoblastic differentiation.

**Figure 2.**
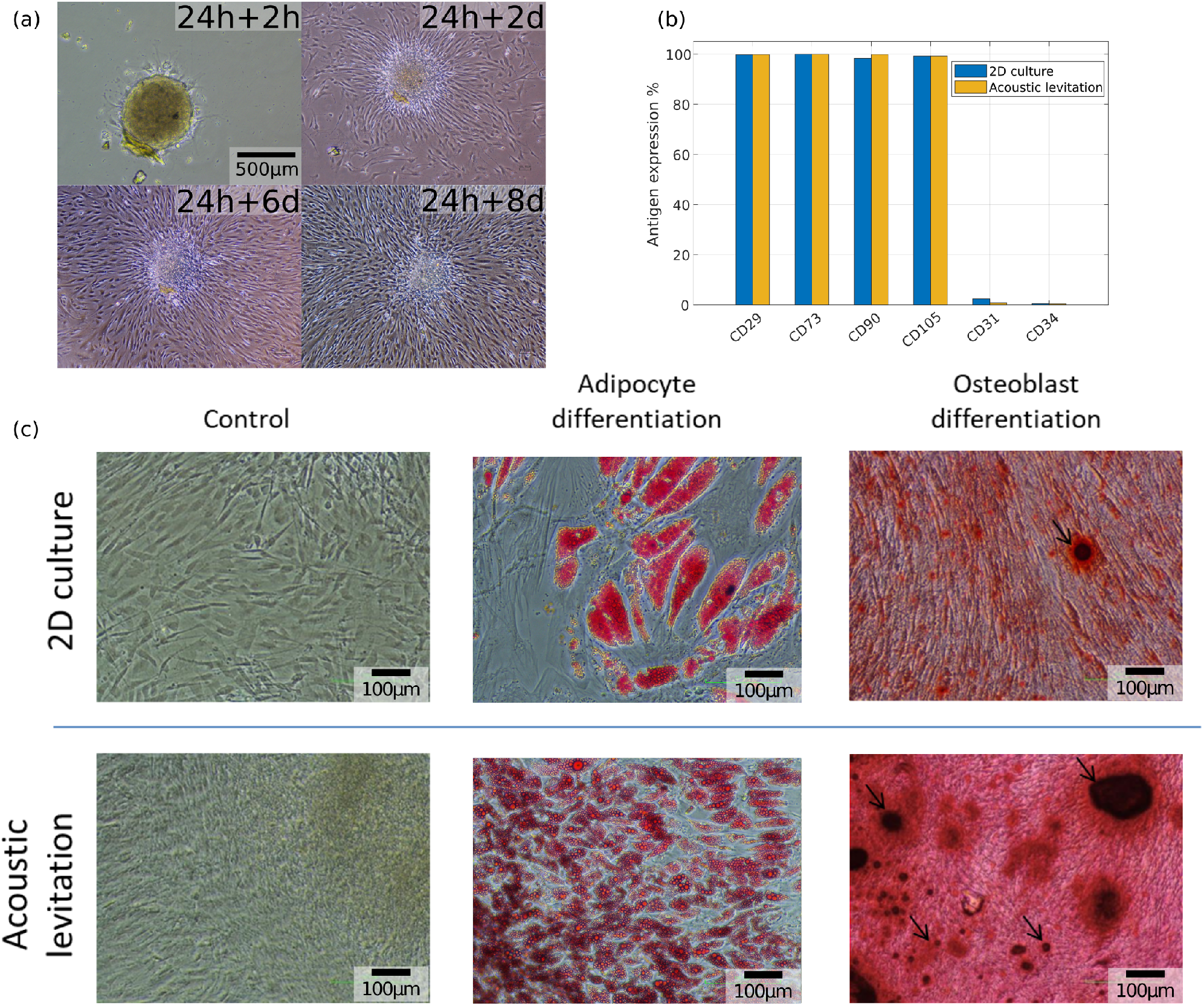
Biological evaluation of MSCs in levitation. (a) Pictures of a spheroid recovered after the 24h levitation and reseeded in a classical culture plate. 2h after the levitation, the spheroid began to adhere to the plastic surface. Then, during the 8 following days, the cells spread out onto the plate. (b) Evaluation of MSC cell surface markers by flow cytometry. After a 24h levitation, the spheroids were expanded in polystyrene dishes. A week later, cells were detached, stained and analyzed for the expression of classic MSC markers by flow cytometry. Here the results are expressed as the percentage of positive cells. (c) Differentiation of MSCs into adipocytes and osteoblasts. After a 24h acoustic levitation, spheroids were reseeded in 24 well plates. Once they adhered, they were treated with MSC expansion medium (control), adipocyte or osteoblast differentiation media. A 2D confluent culture of cells, which did not undergo acoustic levitation was used as a control condition. Adipocyte differentiation was revealed by the staining of lipid vesicles by oil-red O and osteoblasts were revealed by alizarin-red staining.

## Discussion

We hereby demonstrate the development of a robust and reproducible cell culture system based on acoustic levitation using a PDMS chip and an ultrasound transducer, with the possibility of a constant flow, which allows culture medium renewal. Even though previous studies have shown the possibility of using acoustic levitation for cell sheet and aggregate formation, these studies were limited to a very short treatment period usually less than one hour [10]. Only a few studies pushed the culture time beyond 24 hours. However, either the device was limited to a single aggregate [22] or the method produced small aggregates with no levitation (< 150μm) [23]. Thanks to its versatility and ease of implementation, our cell culture in multi-trap acoustic levitation (CCMAL) device allows the production and the culture of hundreds of large spheroids with a reproducible size. The CCMAL setup can be easily scaled up to culture thousands of spheroids in acoustic levitation in parallel. It is well adapted for MSC cultures, preserving their viability, proliferation and characteristic surface markers. It also increases their potential of differentiation. It has been previously shown that high density of MSCs in 2D culture, as well as spheroid formation in low attachment dishes [24], or the application of ultrasound [25] enhances their osteoblastic potential. In our system, the rapid formation of spheroids mimics both high confluency of 2D cultures and spheroid formation owing to the acoustic confinement (aggregation induced by the axial and transverse components of the ARF). This approach opens the path to a better understanding of these phenomena and application to many other different types of cells. It also demonstrates the use of acoustic levitation as a new method of long-term, reproducible cell culture process for scaffold-free tissue engineering purposes such as organoid formation and multicellular co-cultures.

## Materials and methods

### Acoustofluidic setup

For each experiment, we ran four cell culture in multi-trap acoustic levitation (CCMAL) devices in parallel in order to increase the number of spheroids and maximize the cell preparation. One has to control the acoustic levitation process, the optical observation, the fluid perfusion as well as the biological part from the cell preparation to the post-levitation analysis and the collect of all samples.

The ultrasonic waves were generated by a transducer driven by an arbitrary waveform generator (Handyscope HS5 from TiePie Engineering™) monitored by a computer. Each output of the signal generators supplied two ultrasonic transducers (2 MHz SignalProcessing™) with a sinusoidal waveform of amplitude 7Vpp and frequency 2.15MHz. The wave parameters were chosen to optimize the levitation process while avoiding undesired phenomena like acoustic streaming.

The chip is composed by a PDMS (Polydimethylsiloxane) body bonded on microscope cover-glass and a mono-element ultrasonic transducer. The geometry of the PDMS chip is designed to allow the insertion of the ultrasonic transducer inside the PDMS body. The transducer closed the acoustofluidic cavity. One side of the resonant cavity is ultrasound emitter, in contact with the fluid, avoiding acoustic losses through various interfaces, while on the other side the microscope cover-glass acts as a powerful acoustic reflector and allows an optical access. The overall acoustic resonant cavity is simple and efficient.

To shape the PDMS body, we use a negative mold made with a 3D printer (3D Form 2 from Formlabs company). Then the PDMS (RTV 615) is poured in and put in the oven at 70°C for 24h. After unmolding the solid PDMS body is bounded on a cover-glass with a plasma cleaner.

Because of this fabrication process, the geometry of the chip is highly versatile. We settled on two geometries. The first chip (see **Figure 3a**) includes the levitation cavity with 11 nodes, a channel and a simple bubble trap. Our manufacture process makes the sidewall perfectly transparent to visible optical waves and allows the observation from the side of the acoustic levitation process over the entire cavity. To increase the spheroid number from 11 up to 27 we designed a second chip.

**Figure 3.**
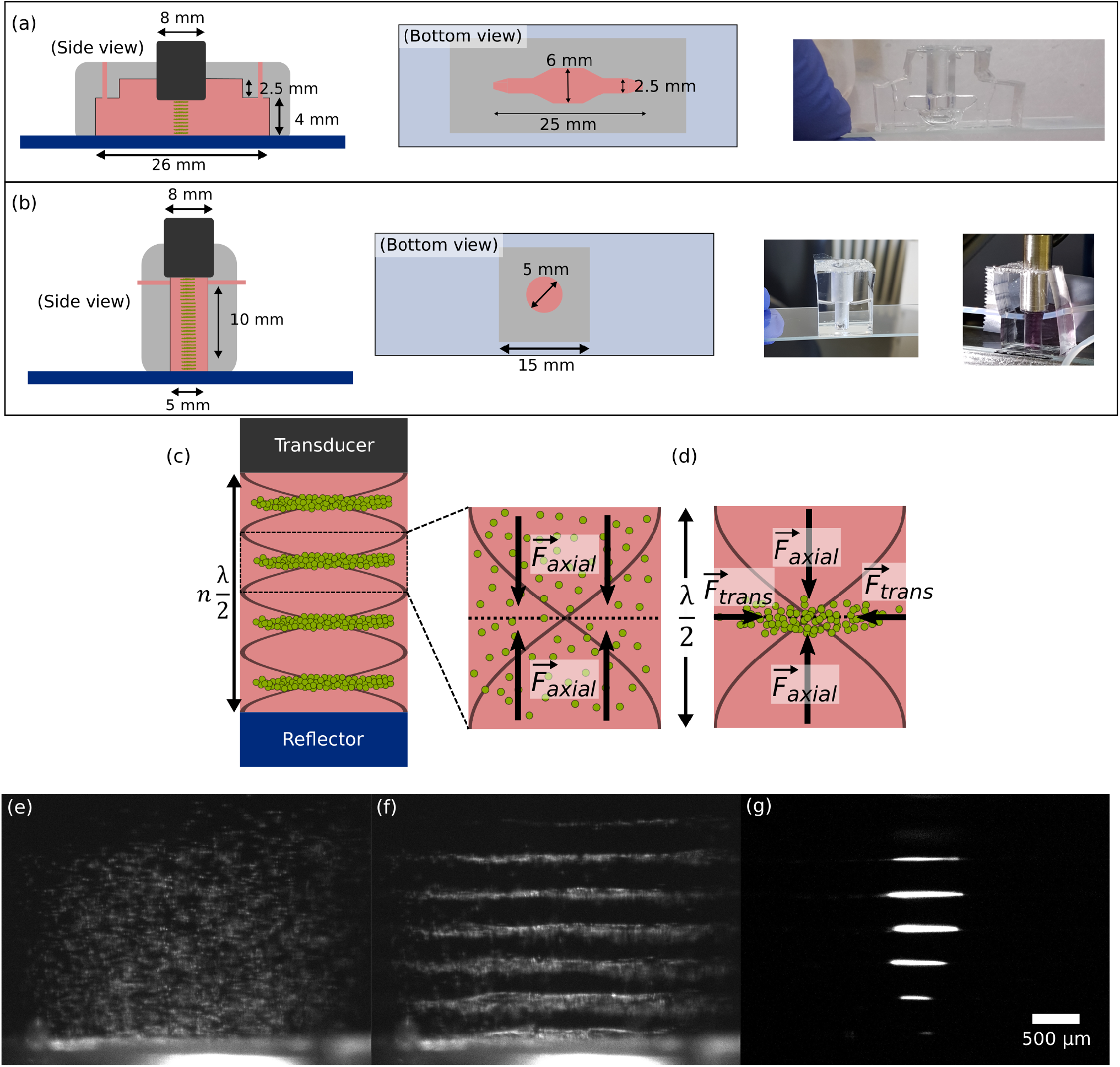
(a) Chip geometry 1 (b) Chip geometry 2 (c) Schema of the multilayer acoustic cavity (d) The suspension of cells particles undergoes the Acoustic Radiation Force (ARF). The cells are first focused in the nodal plane and then, trapped by the transverse component of the ARF. The cavity dimensions are adjusted to have multiple clusters. Example of the levitation and aggregation process: (e) Suspended particles are injected in the cavity. Here is 15μm diameter polystyrene beads. (f) Axial acoustic radiation forces are dominating and bring the particles in the pressure nodal plane. (g) Therefore, the transverse forces trap particles in a circular monolayer.

The geometry (see Figure 3b) consists in a cylindrical well of diameter *D* = 5mm and height 10mm. The optical access is still possible but with a lower quality. To monitor the formation of the cells sheets and the spheroids self-organization from the side view, we used four usb video microscopes (Edge and AM 2111 from Dinolite company).

The culture medium was continuously perfused in the chip with a syringe pump (Pump 11 Elite from Harvard Apparatus™) in a withdrawal mode with a flowrate of 20μL/h small enough to avoid any perturbations inside the acoustofluidic cavity.

### Image processing for the study of the spheroid size

In order to characterize the time evolution of the sizes of the spheroids, we developed a Matlab image analysis code. It works automatically with a few user-defined parameters. In particular, we manually defined the regions of interest (ROIs) for each spheroid. Then an ellipsoid is fitted on the binarized image. The section of the original image passing through the ellipse center is selected in the axial and radial directions. We stored the width of the 70% thresholded curves of the X and Y sections (See movieImageProcessing.mp4). For the whole image sequence, ROIs were defined automatically from the ellipsoid characteristics of the previous image. We obtained the time evolution of the axial and radial dimensions of the spheroid with a one-minute time step. We finally applied a moving average of 30 points to smooth the signal fluctuations. Afterwards, the average and the standard-deviation were computed.

### MSC isolation and characterization

MSCs were isolated from adipose tissue of the thigh of a healthy donor, after a signed consent, according to the French regulations. MSCs were obtained after the digestion of adipose tissue with collagenase NB6 (Serva electrophoresis). The adipose tissue was placed in 40ml of collagenase NB6 diluted in α-MEM medium at a final concentration of 5μg/ml in a 50ml conical tube (Falcon, Dutscher). After a 2 hour incubation at 37°C, the digested tissue was filtered through a cell strain with pores of 100μm to remove the remaining tissue. The strained cells were then centrifuged and plated in a cell factory (Thermofisher™, Nunc™) for expanding in MSC culture medium composed of α-MEM (Gibco™) supplemented with 10% fetal bovine serum (FBS, Biowest) and 1% antibiotic / antimycotic mix (Anti-Anti 100x, Gibco™).

At the second passage, cells were characterized by flow cytometry using a panel of MSC positive markers including CD29-PE (MACS Miltenyi), CD73-PE (BD), CD90-FITC (BD), CD105-PE (BD), and negative markers including CD31-FITC (BD) and CD34-APC (BD). Unstained cells and cells stained with control isotypes have been used as negative control.

### MSC characterization after acoustic levitation

After 24h of acoustic levitation, the spheroids were collected and immediately fixed overnight in paraformaldehyde 4% diluted in PBS. They were then embedded in a fibrin gel to facilitate manipulation for preparing samples. The fibrin gel containing the spheroids was included in paraffin and 5μm thick sections were cut. The slices were stained with H&E for general structure. Ki67 immunohistochemistry was performed with anti-ki67 antibody (Agilent-Dako) 3-step immunoperoxidase technique.

### MSC viability and adherence after acoustic levitation

To evaluate the capacity of MSC spheroids, which also shows their viability, they were placed in 24 well dishes and were observed during 24h in an Incuyte. Cell viability was further evaluated with a LIVE/DEAD™Viability/Cytotoxicity Kit (Invitrogen). The components (calcein and propidium iodide) were diluted in PBS according to the provided instructions of the kit and the spheroids were incubated in the solution for 30 mins in an Ibidi™dish prior to observation with a confocal microscope (Zeiss LSM 780 AiryScan).

### Evaluation of MSC surface markers and potential of differentiation

After 24h of acoustic levitation, MSC spheroids were collected and reseeded in 75cm^2^ for expanding in MSC culture medium. About a week later, cells were detached and evaluated by flow cytometry using a panel of MSC positive and negative markers including CD31, CD34, CD29, CD73, CD90 and CD105. Data were acquired and analyzed with an Attune flow cytometer (Thermofisher™).

The potential of differentiation of the aggregates into adipocytes and osteoblasts was also evaluated. For this purpose, the spheroids were collected and reseeded in a 24 wells plate at a density of one spheroid / well in MSC expansion medium. A 2D culture of cells at the same passage that had not undergone levitation was also used as control.

After allowing the cells to adhere and expand for three days, the cell culture medium was replaced by adipocyte differentiation medium or osteoblast differentiation medium (StemX-Vivo™, R&D systems). MSC expansion medium was also used as the control condition. After a two-weeks culture, cells were fixed with 4% paraformaldehyde. Adipocyte differentiation was permeabilized with isopropanol for 5 mins and the lipid vesicles were stained with oil red-O (Sigma-Aldrich). Osteoblasts were stained with alizarin red (Sigma-Aldrich).

## Supporting information

movieBeads

movieCellLevitationEx1

movieCellLevitationEx2

moviePh

movieGFP

movieImageProcessing

## Ethical statement

We declare that all methods in the manuscript were carried out in accordance with relevant guidelines and regulations given by Advanced Healthcare Biomaterials. All procedures involving patients were conduct according to the Helsinki Declaration.

Mesenchymal Stromal Cells were isolated from the adipose tissue of a male donor, obtained at Necker-Enfants Malades hospital in Paris, France. The adipose tissue was the surgical leftover and was used for research purposes after a signed consent from the donor and his parents as the legal tutors, according to the French bioethical and medical research regulations. In France, according to the law “loi Jardé” (article L. 1121-1 of the public health code) governing scientific research on human subjects and tissue samples, the surgical leftover can be used for scientific research without the prior approval of an ethical committee.

## Acknowledgements

The authors would like to thank Briac Thierry and Françoise Remangeon for the collect and the donation of MSCs, and especially Briac for his support on the ethical rules.

## Author’s Contributions

N.J. and L.A designed and performed experiments and wrote the manuscript, N.S. designed and performed experiments, P.B. performed the immunohistochemistry analysis, M.H. supervised the project, J.L. and J-L.A. conceived and supervised the project, and reviewed the manuscript.

## Supplementary informations

### Acoustofluidic principles

The acoustic levitation describes the fact that an object is suspended in a medium by the action of the Acoustic Radiation Force (ARF) against the gravity. The ARF results from the time-averaged acoustic quantities from scattering of the acoustic waves on the objects. The ARF is explained by the second order perturbation theory of the Navier-stokes equation. The first analytical models of the ARF applied to particles date back in 1934 and make strong simplifying hypothesis like considering incompressible spheres [26]. Then, the compressibility was taken into account by the Yosioka model [27] but with sound field assumptions (considering a plane progressive or a standing-plane wave). Gor’kov [28] summarized and generalized the previous models with the following assumptions: the surrounding fluid is inviscid and the particles are much smaller than the acoustic wavelength λ.

In our experiments we used the axial and the transverse components of the ARF: the axial component moves the cells toward the pressure nodes, while the transverse components forces the cells to gather at the location of maximum acoustic energy, roughly at the center of the cylindrical cavity. The location and strength of the acoustic traps can be adjusted by changing the acoustic frequency *f*_*ac*_, as shown by [29]. The acoustic levitation process is mainly due to the primary ARF given by the equation (1).

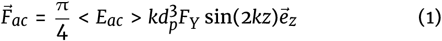

with < *E*_*ac*_ > for time-average of the acoustic energy density, 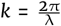 the wave number of the acoustic plane wave with frequency *f*, *c*_*m*_ the acoustic celerity in the medium, *d*_*p*_ the particle or cell diameter, and *F*_*Y*_ the contrast factor. The *z* variable represents the axial position. The cells are affected by the primary ARF because of their acoustic contrast with the suspending medium quantified by the acoustic contrast factor *F*_*Y*_ :

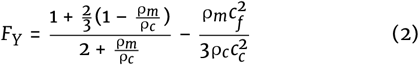

with *c*_*p*_ the acoustic celerity in the cell material, *c*_*m*_ the acoustic celerity in the medium, ρ_*c*_ the cell density and ρ_*m*_ the medium density.

The amplitude of the primary ARF typically ranges between 10^−14^ and 10^−12^ N (here for a 5Vpp input signal, the force is about 5 × 10^−11^ N).

In general, for cells and micro-organisms like bacteria, the contrast factor is positive thus they move toward the pressure nodes. The closer the cells go near the theoretical node, and the weaker the axial primary ARF will be until cancelling itself when reaching the pressure node. From this observation, the transverse ARF becomes significant and is responsible of the cell aggregation in the nodal plane. For a radially symmetric acoustic wave, the transverse ARF can be described in the nodal plane with the equation equation (3)

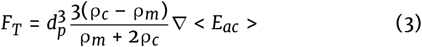

In this work we use a cylindrical transducer. The radial acoustic energy gradient is directed toward the axis center, where the acoustic energy is the highest. This region can be considered as an acoustic trap where any cells with a positive contrast factor, as well as most living micro-organisms, even motile like bacteria [30], can be trapped and contained within the acoustic force field [31].

A supplementary short-range force, called Bjerkness or Secondary Radiation Force (SRF), attracts cells close to each other until clustering. This is an inter-cells force due to the scattered acoustic field by the other cells. When cells are lined up perpendicular to the wave propagation (located in the nodal plane for instance) the force is attractive and can be expressed by equation (4)

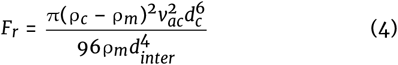

with *v*_*ac*_ the acoustic velocity at the plane position of the cells and *d*_*inter*_ the distance inter-cells.

The combination of these forces allows the creation of layers of cells at every pressure node. In this work we used a robust approach to obtain efficient levitation and aggregation, which consists in generating pseudo standing waves in a resonant cavity. Indeed, to maximize the emitted energy, the acoustic wave is confined between the transducer and the reflector. To obtain constructive interferences between the emitted and the reflected wave, the height *H* of the cavity must be equal to a multiple of the half-wavelength 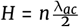.

## Note

The article layout was adapted from the latex template for GigaScience Journal Manuscript Soumissions under the Creative Commons CC BY 4.0. The authors of this template are OUP and Overleaf. https://www.overleaf.com/latex/templates/template-for-gigascience-journal-manuscript-submissions/shgtrssvbjhs

